# Delta PSA: A New Metric for Conformational Dynamics Underlying Macrocyclic Peptide Permeability

**DOI:** 10.64898/2026.01.06.697862

**Authors:** Yiping Yu, Qilong Wu, Junwei Chen, Yuli She, Lizhe Zhu, Zhiye Guo

## Abstract

Development of oral cyclic drugs often suffers from low oral bioavailability resulting from limited passive permeability making accurate prediction a central challenge in drug development. Existing approaches generally fall into two categories: deep learning–based models and accelerated molecular dynamics (aMD) simulations. While deep learning models enable rapid, high-throughput predictions, they often suffer from dependence on training data, and sensitivity to dataset biases. On the other hand, aMD provide mechanistic interpretability by explicitly modeling peptide translocation across lipid bilayers, however, suffer from huge computational cost.

Here, we present a conventional MD-based framework for predicting macrocyclic peptide permeability, designed to facilitate interpretation. Importantly, we show that **Delta PSA** (Δ*PSA*) directly quantifies a peptide’s chameleon propensity, providing a mechanistically meaningful measure. Furthermore, we identify two key structural indicators—the sidechain PSA ratio and the radius of gyration—as refined metrics for assessing conformational behavior. When applied to a benchmark dataset of macrocyclic peptides, our framework achieves an MSE of 0.22, surpassing the 0.25 reported for Multi_CycGT and demonstrating its superior performance. Our work also provides a large dataset of MD trajectories for macrocyclic peptides in both polar and nonpolar environments. This dataset offers access to a broader conformational space for cyclic peptide studies.

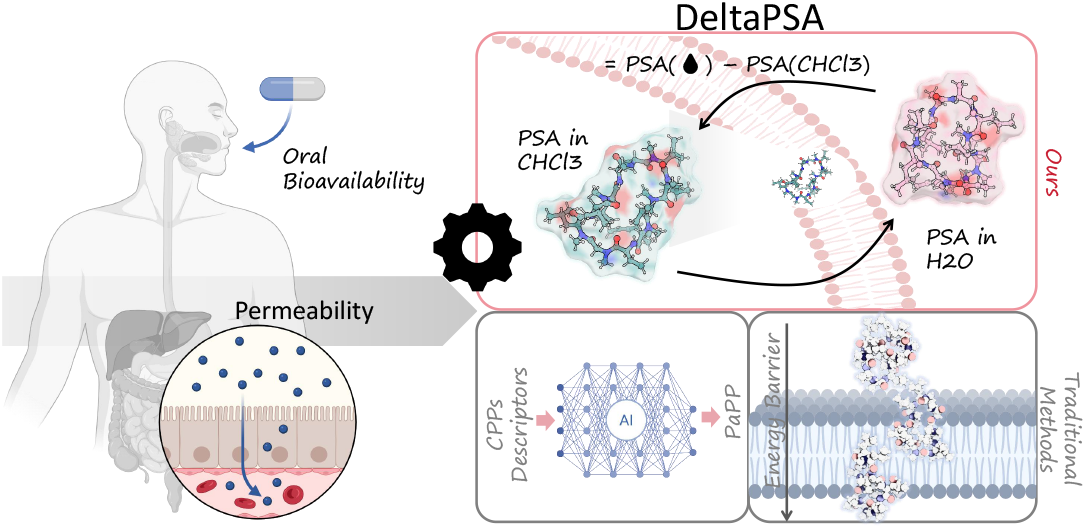

## 1. Introduction

The clinical success of cyclic peptide drugs is mounting, with over 40 currently in use and the FDA’s approval of roughly one macrocyclic peptide annually for the past 20 years [36, 47, 48]. Cyclic peptides, which combine the precision of biologics with the developmental advantages of small molecules, are attracting growing interest. The balance of specificity, stability, and synthetic accessibility makes them powerful modalities for targeting intractable diseases.

Despite their pharmacological potential, the druggability of cyclic peptides has emerged as a critical bottleneck in the development of next-generation peptide therapeutics. Cyclic peptides often possess superior metabolic stability and binding specificity compared to their linear counterparts. However, their inherently poor membrane permeability poses a major challenge to their clinical translation [29, 42]. To traverse lipid bilayers, macrocyclic peptides typically exploit conformational adaptability and physicochemical properties that enable passive membrane permeation without compromising aqueous solubility [1]. Therefore, a mechanistic understanding of these processes is essential for the design of cyclic peptides that target intracellular proteins.

While all peptides exhibit some conformational flexibility, the chameleonic behavior of certain cyclic peptides—where they can adopt multiple, structurally diverse, nearly isoenergetic conformations—is particularly pronounced and functionally critical. This conformational plasticity reconciles two opposing demands: in polar environments the peptide can expose polar groups and hydrogen-bond with solvent to maintain solubility, while in nonpolar or membrane-like environments it can reorganize to form intramolecular hydrogen bonds (IMHBs) that shield polar atoms, lower desolvation penalties, and favor passive membrane insertion. Prototypical examples such as Cyclosporin A illustrate how backbone N-methylation and internal hydrogen bonding enable oral bioavailability despite violations of classic small-molecule rules [1, 13, 25]. This environment-dependent reorganization not only facilitates membrane transition by decreasing polar surface exposure in nonpolar media but also preserves the peptide’s ability to present binding determinants in aqueous environments—thereby maintaining high target affinity [20, 32, 45]. Structural and biophysical studies (e.g., NMR, crystallography) have provided direct evidence for such conformational switching and support its exploitation in rational drug design [33].

Computationally, a broad spectrum of approaches has been developed to predict peptide membrane permeability, broadly categorized as molecular dynamics (MD) simulations, quantitative structure–activity relationship (QSAR) models, and machine learning (ML)–based frameworks including recent deep learning implementations [34]. MD simulations yield atomistic descriptions of conformational dynamics and peptide–lipid interaction, advanced sampling techniques (replica exchange MD (REST), steered MD, replica-exchange umbrella sampling (REUS), etc.) have been used to extract 3D descriptors that characterize IMHB patterns, dehydration in the membrane core, and energetic barriers to translocation [19, 39, 41]. In some cases, the energy barriers revealed by aMD simulations with explicit cholesterol-containing bilayers have shown high correlation with experimental permeability (*R >* 0.8) [38]. However, simulating complete transitions with aMD remains computationally intensive and often fails to adequately sample rare, yet functionally critical, conformations. This challenge is further exacerbated by slow processes like the cis/trans isomerization of N-methylated or proline residues, which introduce high energy barriers and significant conformational heterogeneity [11, 26].

Quantitative structure-activity relationship (QSAR) models offer efficient alternatives by correlating key molecular descriptors—such as logP, polar surface area, and hydrogen bond donor/acceptor counts—with permeability. Some advanced variants go beyond simple 2D parameters by incorporating 3D descriptors that can represent molecular flexibility and intramolecular hydrogen bonds (IMHBs) [19, 34]. Yet QSAR approaches are inherently limited by their reliance on static features and often fail to capture the environment-dependent chameleonic behavior of macrocycles.

To address these shortcomings, ML and deep learning frameworks have been developed that integrate multi-level molecular representations. Conventional ML algorithms (Random Forests, SVM, XGBoost) show moderate performance [18, 40], while multimodal deep architectures—such as frameworks combining atom-level, monomer-level and peptide-level features or fusing graph convolutional networks with Transformer encoders—have achieved stronger predictive metrics (e.g., CycPeptMP: *RMSE* ≈ 0.503, *R* ≈ 0.883; Multi_CycGT: *accuracy* ≈ 0.8206, *F* 1 0.8708) [6, 18]. Hybrid methods that incorporate MD-derived dynamic descriptors into ML pipelines also show promise by blending mechanistic insight with statistical learning [39]. Despite these advances, a significant gap remains: these models primarily operate on static molecular representations or single-conformer inputs and are not explicitly designed to learn from the dynamic conformational ensembles that peptides sample in different environments.

The Rule of Five highlights molecular weight (MW) and polar surface area (PSA) as critical for passive permeability. We confirmed this for lower-MW cyclic peptides (*<* 800 Da), where permeability negatively correlates with topological polar surface area (TPSA), as validated by the CycPeptMPDB database (**Fig. S1a**). However, many larger cyclic peptides (MW *>* 800 Da) defy this rule and exhibit unexpected permeability, which static models cannot explain [12, 24, 31].

This gap is particularly critical for chameleonic peptides, which exist not as single conformers but as dynamic ensembles of rapidly interconverting states, yet they are frequently underrepresented in standard computational sampling [7]. While conventional logP and fragment-based lipophilicity calculations typically overlook the critical roles of dynamic conformations [2, 21], this limitation is exacerbated in larger, more flexible macrocycles. Duan’s lab tested Multi_CycGT on peptides of 6–7-residue and other lengths, finding that all metrics decreased for the non 6–7-residue peptides (dominated by 10-residue peptides) [40].

These limitations underscore the need for methods that can simultaneously account for dynamic conformational flexibility and solvent-dependent effects [14, 27, 34, 43]. We thus propose a ‘conformational switching’ model. It posits that large cyclic peptides are dynamic, toggling between lipophilic (membrane-buried) and hydrophilic (solution-exposed) states. We hypothesize that this conformational adaptability, not static descriptors, dictates their transmembrane potential. To test this, we selected cyclic peptides with ≥ 9 amino acids (MW *>* 800 Da) as an ideal model system to investigate this dynamics-driven permeability (see in **Fig. S1b**). This study aims to elucidate the membrane permeation mechanisms of macrocyclic peptides.

In response, our work leverages conformational ensembles sampled from conventional molecular dynamics (cMD) in contrasting solvents (water and chloroform). We posit that MD trajectory-derived dynamic features serve as superior indicators by directly reflecting the conformational adjustments inherent to chameleonicity. For instance, solvent-accessible surface area (SASA) occupancy profiles quantify how pharmacophore groups are dynamically exposed or shielded during transitions between aqueous and nonpolar environments [15, 17, 28]. Reductions in PSA frequently correspond to “closed” conformations favorable for bilayer insertion [34]. Crucially, long-range hydrogen bonds and decreases in PSA observed in nonpolar states are mechanistic hallmarks of chameleonic switching—properties often overlooked by static descriptors and may be inadequately sampled in conventional MD simulations [44].

SASA, in particular, gives a solvent-specific snapshot but does not intrinsically quantify the magnitude of the environmental adaptation that is central to the chameleonic effect. To address this gap, we introduce the difference in polar surface area (ΔPSA) between aqueous and membrane-mimicking environments. ΔPSA is a dynamic feature extracted directly from comparative MD trajectories, it explicitly measures the reduction in exposed polarity as a peptide reorganizes from a hydrophilic to a hydrophobic phase, thereby capturing the desolvation advantage conferred by conformational dynamics that underpins passive membrane permeation. As illustrated in **Fig. 1**, macrocyclic peptide like Cyclosporin A undergoes a pronounced PSA decrease when moving from aqueous to chloroform environments, directly linking ΔPSA to its ability to achieve both solubility and permeability through conformational switching. This dynamic property, ΔPSA, along with other trajectory-derived features such as IMHB patterns, allows us to directly link a peptide’s conformational switching capability to its ability to reconcile solubility with permeability.

**Figure 1.**
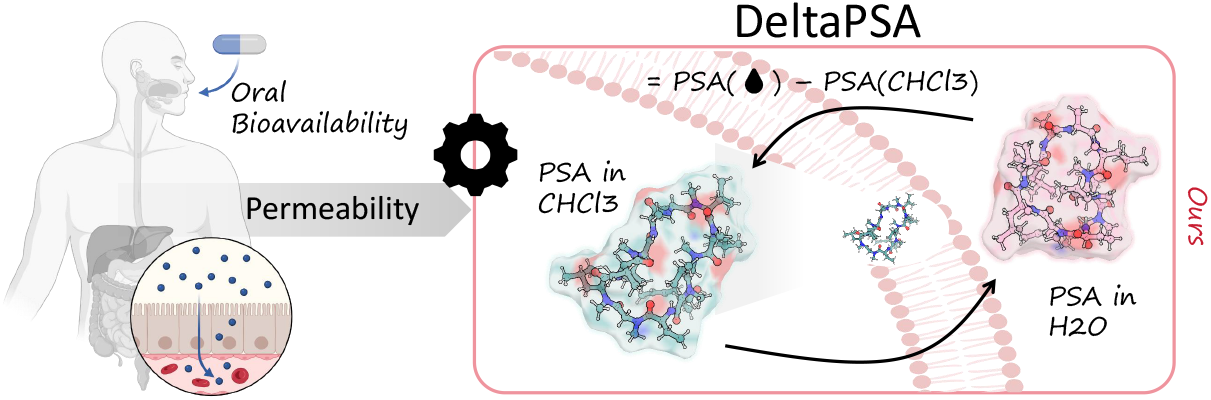
Schematic of the ΔPSA-based method for assessing macrocyclic peptide membrane permeability. Cyclosporin A is shown in stick representation, these conformations were derived from MD simulations in water (colored in red) and chloroform (colored in green), respectively.

Building on these insights, the present study develops a cMD-based framework that leverages cMD trajectories to extract SASA, ΔPSA, and related dynamic descriptors for predicting cyclic peptide permeability using evaluating chameleonic propensity. By simplifying the MD workflow for macrocyclic peptides, our approach reduces computational cost while retaining mechanistic interpretability. Importantly, focusing on SASA and ΔPSA dynamics enables us to (i) identify permeable peptides with chameleonic propensity, (ii) relate interpretable descriptors to permeability, and (iii) provide a practical surrogate for computationally expensive enhanced sampling methods. Applied to a benchmark dataset, this framework achieves an MSE of 0.21 for permeability prediction, demonstrating that dynamic MD-derived features can deliver both predictive accuracy and mechanistic insight for macrocyclic peptides.

## 2. WorkFlow for Δ*PSA* Method

### 2.1 Conformational Ensemble Extracted from MD Sampling

Several studies have focused on developing conformational sampling methods for cyclic peptides and macrocycles. Li et al. generated 3D conformations of peptides and their monomers using the RDKit package (version 2022.09.5), where initial structures were constructed via the ETKDG method in vaccum and subsequently optimized through energy minimization with the UFF force field [18].

While these approaches improve the generation of initial conformations in vacuum, they are still limited in exhaustively capturing the conformational diversity of macrocyclic peptides, especially for those exhibiting chameleonic behavior. Accelerated molecular dynamics (aMD) sampling has therefore been explored as an alternative, providing deeper sampling of conformational ensembles [9]. However, advanced enhanced-sampling methods such as metadynamics and adaptive biasing approaches come with considerable computational costs [16, 22], and are not always practical for large-scale peptide screening. Conventional MD (cMD) remains widely used due to its simplicity and physical interpretability, but its sampling efficiency is often insufficient to explore rare conformational transitions within feasible timescales [9].

In our pipeline (see Fig. 2 a), MD simulations were carried out for macrocyclic peptides ( ≥9 residues; CycPeptMPDB) using a small-molecule representation. Initial structures were retrieved directly from CycPeptMPDB entries in water. All simulations were conducted using the Gromacs package (version 2023.3), with force field parameters generated by the Sobtop software and the TIP3P water model [3].

**Figure 2.**
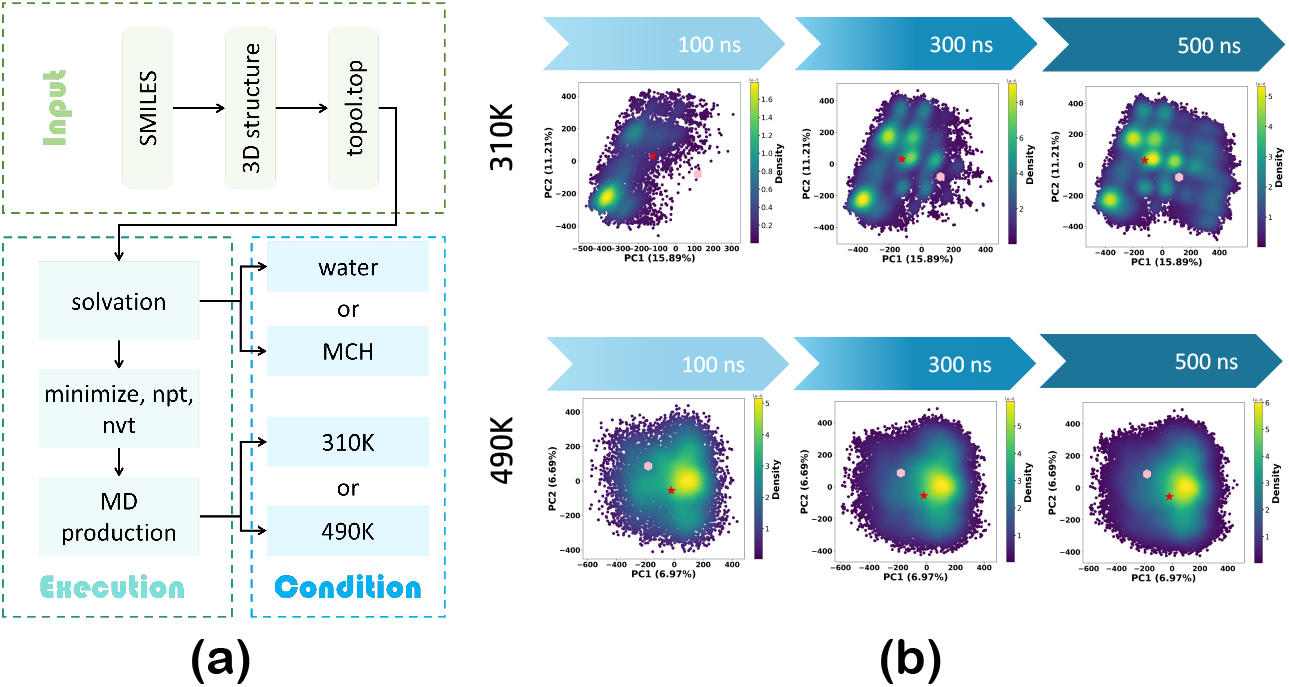
**(a)** Workflow for production MD of a macrocyclic peptide. **(b)** PCA projection density from ring dihedral angles at 310 K and 490 K.

Simulation systems were constructed using the Gromacs gmx solvate module to efficiently solvate the cyclic peptides in either water or chloroform boxes. The size of each solvent box was chosen to maintain a minimum distance of 1.0 nm between the peptide and the box boundaries, preventing artificial interactions between periodic images. Counterions were added using gmx genion to neutralize the system, ensuring overall charge neutrality and simulation stability.

Prior to production runs, energy minimization was performed using the steepest descent algorithm for up to 50,000 steps to remove steric clashes or unfavorable van der Waals interactions. Following minimization, a two-stage equilibration procedure was applied. The first stage was a 100 ps NVT equilibration (constant number of particles, volume, and temperature) using the V-rescale thermostat with a coupling time constant of 0.1 ps to gradually bring the system to the target temperature [35]. The second stage was a 100 ps NPT equilibration (constant number of particles, pressure, and temperature) with the Parrinello-Rahman barostat and a coupling time constant of 2.0 ps to adjust the system to 1 bar, ensuring correct solvent density [46].

Production simulations were run for 500 ns with a 2 fs time step, saving coordinates every 10 ps for analysis. The LINCS algorithm constrained all bond lengths involving hydrogen atoms, allowing the use of a larger time step without compromising accuracy [9]. Long-range electrostatic interactions were treated using the Particle Mesh Ewald (PME) method with a real-space cutoff of 1.2 nm [10]. Van der Waals interactions were truncated at 1.2 nm, applying a force-switching function between 1.0 nm and 1.2 nm for smooth energy transitions.

To explore conformational differences in lipid-like and aqueous environments, peptides were simulated separately in chloroform and water boxes. Both normal-temperature (310 K) and high-temperature (490 K) simulations were performed to assess the impact of temperature on conformational sampling. The elevated temperature was intended to accelerate rare conformational transitions that might be inaccessible at physiological temperature.

Notably, conventional MD at physiological temperature often suffers from limited sampling, especially for macrocyclic and chameleonic peptides whose conformational landscapes include high-energy states that are critical for membrane crossing. Existing conformer generation methods, such as ETKDG or Prime-MCS, provide reasonable starting structures but are generally insufficient for exhaustive sampling of environment-dependent chameleonic states [35, 46]. By employing high-temperature MD, we can overcome kinetic barriers, more efficiently explore diverse conformational states, and capture transitions between aqueous-favored and apolar-favored structures. PCA convergence analysis were used to validate the sampling adequacy (see **Fig. 2b**).

#### Convergence Analysis Validates Molecular Dynamics Sampling

We assessed MD sampling convergence by examining 500-ns trajectories of Cyclosporin A (a typical chameleonic peptide) in aqueous solution at temperatures of 310 K and 490 K. For each trajectory, the dihedral angles of all ring atoms were calculated and subjected to principal component analysis (PCA), with the first two principal components used for visualization in a two-dimensional (2D) space (**Fig. 2b** ) [3].

To quantitatively evaluate convergence, we compared the 2D density distributions of trajectory segments of varying lengths (100, 300, 500 ns). At 310 K, the system did not reach convergence until 500 ns, in contrast to the much earlier convergence observed at 490 K. These results are consistent with the expectation that elevated temperature accelerates barrier crossing and enhances conformational sampling.

Taken together, the PCA density plots and density-difference analysis (**Fig. 2**) confirm that 500 ns simulations are sufficient to reach equilibrium at physiological temperature, while high-temperature MD substantially improves sampling efficiency. The statistical metrics derived from these equilibrated trajectories can thus be regarded as reliable approximations of the true conformational distributions under their respective conditions.

### 2.2 Dynamic Featurization based MD Trajectories

To capture the structural and dynamic characteristics of cyclic peptides from molecular dynamics (MD) simulations, we developed a comprehensive feature extraction workflow that integrates MDAnalysis with GROMACS utilities. This workflow systematically computes a wide range of features—including conformational properties, flexibility, solvent accessibility, and intramolecular interactions—that are critical for downstream model training and analysis.

Conformational and flexibility features (**Table 1**) include RMSD with statistical summaries, RMSF to identify dynamic flexibility, and analyses of key bond angles. Solvent accessibility and spatial properties are quantified through SASA and radius of gyration to assess compactness and mass distribution. Intramolecular interactions are evaluated through hydrogen bond analysis to track counts, variability, and stability, along with specialized monitoring of disulfide bonds and cyclic motifs.

**Table 1.** Structural and Dynamical Properties of the Conformational Ensemble.

| Descriptor | All atoms | Backbone | Sidechain | Polar | Nonpolar |
| --- | --- | --- | --- | --- | --- |
| RMSD | ✓ | ✓ |  |  |  |
| RMSF | ✓ | ✓ | ✓ |  |  |
| Bond angles |  | ✓ | ✓ |  |  |
| SASA | ✓ | ✓ | ✓ | ✓ | ✓ |
| Gyration | ✓ | ✓ | ✓ |  |  |
| Hydrogen Bond | ✓ |  |  |  |  |
| Disulfide Bond | ✓ |  |  |  |  |

All extracted features are aggregated and exported as structured data suitable for statistical analysis or machine learning. By capturing critical dynamic and structural characteristics, this approach provides robust input for predictive modeling of cyclic peptide properties of membrane permeability.

## 3. Results

### 3.1 Data Distribution in Δ*PSA*

Our analysis of the CycPeptMPDB database revealed the length distribution and permeability characteristics of the macrocyclic peptides. The peptide lengths span from 9 to 15 residues, with a predominance of 10-residue peptides (**Fig. 3a**). Furthermore, the ratio of permeable (logP *>* −6) to nonpermeable (logP ≤−6) peptides was assessed as a function of length (**Fig. 3b**). Notably, the 10-residue peptides show a ratio of 1.69, indicating a state of relative balance between the two permeability groups.

**Figure 3.**
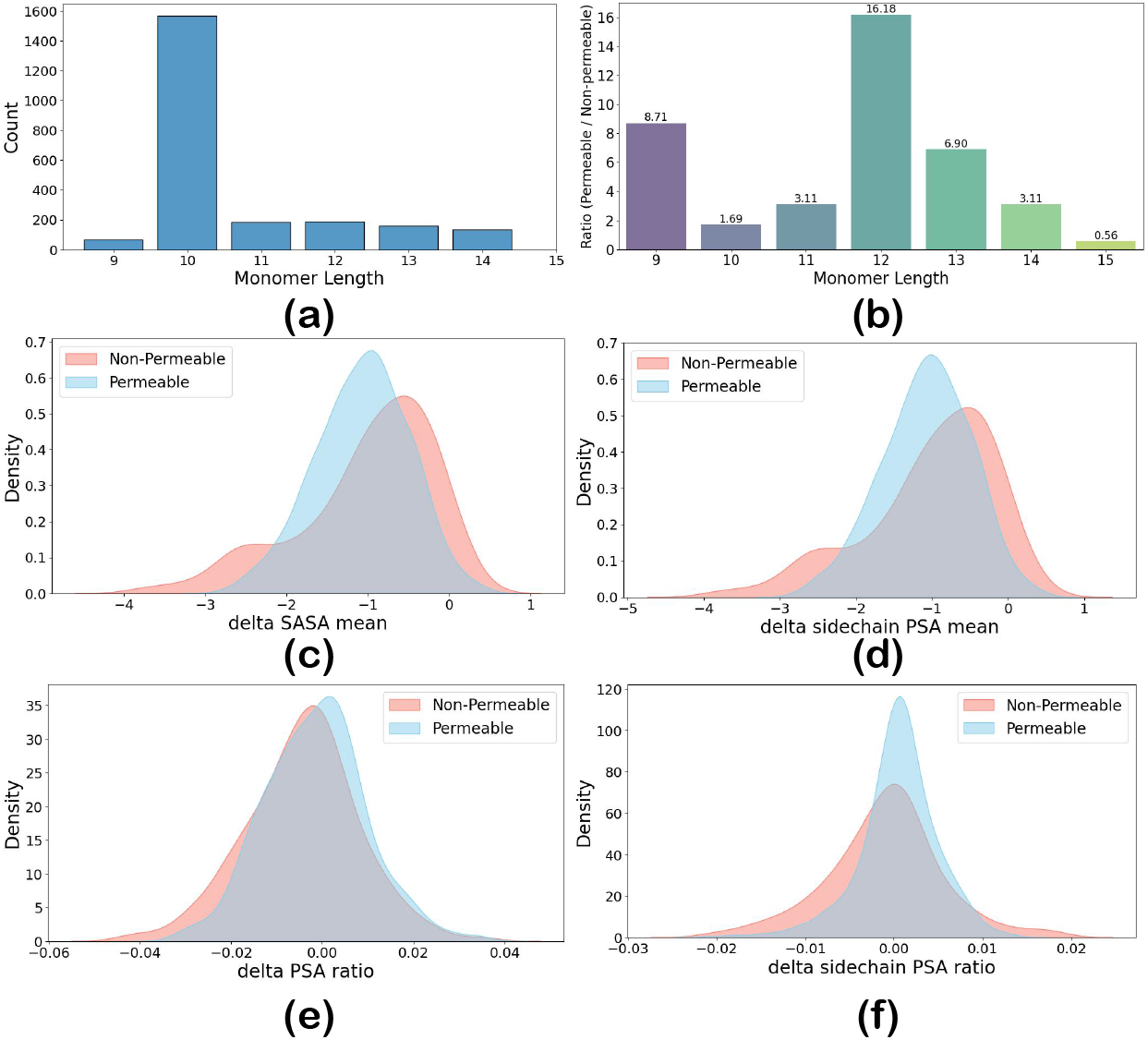
**(a)** Distribution of Monomer Length in database. **(b)** Ratio of 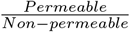 peptides in range of Monomer Length. **(c)** Distribution of 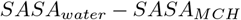 between permeable and non-permeable peptides. **(d)** Distribution of *sidechain* 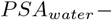 *sidechain PSA*_*MCH*_ between permeable and non-permeable peptides. **(e)** Distribution of 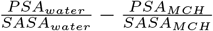 between permeable and non-permeable peptides. **(f)** Distribution of 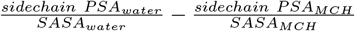 between permeable and non-permeablepeptides.

#### Permeable Macrocyclic Peptides Exhibit Higher PSA Ratio Than MCH in Water

The conformational adaptation of macrocyclic peptides with differing permeabilities was probed by computing the differential SASA (Δ*SASA*^*∗*^ = *SASA*^*∗*^*water* - *SASA*^*∗*^*MCH* ). The observed increase in SASA for permeable macrocyclic peptides within the MCH environment indicates enhanced ring flexibility and a tendency to adopt more open conformations in a hydrophobic milieu. This conformational adaptability is likely a critical factor facilitating membrane permeation (**Fig. 3c,d**). Conversely, the lack of a consistent trend in the Δ*SASA* of non-permeable peptides points to conformational rigidity or instability. This distinct behavior is further mirrored in the Δ *sidechain SASA*, reinforcing the link between peptide flexibility, hydrophobic stabilization, and permeability. Calculation of the Δ*PSA ratio* (PSA/SASA) revealed that permeable macrocyclic peptides predominantly exhibited positive values for both backbone and sidechain ratios (**Fig. 3e,f** ). In contrast, the ratios for non-permeable peptides were distributed around or below zero. This divergence suggests that the permeable peptides form more intramolecular hydrogen bonds (IMHBs) during membrane permeation. A more detailed classification of the delta SASA can be found in the Supporting Information (**Fig. S2**).

In conclusion, permeable peptides exhibited a more pronounced reduction in both total and sidechain SASA, coupled with a larger increase in PSA ratio. This suggests that in an apolar environment, these peptides adopt a more extended conformation (evidenced by the larger absolute SASA in MCH) while simultaneously shielding their polar groups (evidenced by the increased PSA ratio). The marked Δ*PSA ratio* in permeable peptides underscores a superior chameleonic ability, crucial for effective membrane crossing.

To further validate this hypothesis and characterize the underlying conformational landscape, we performed clustering analyses on 1 *µ*s combined MD trajectories for each cyclic peptide (500 ns in aqueous solvent and 500 ns in chloroform at 310 K, see in **Fig. 4**). One frame was extracted every 1 ns, yielding 1000 frames per peptide. Backbone RMSD was calculated for all pairs of frames to construct a 1000×1000 RMSD matrix, and each row of this matrix was treated as a feature vector for KMeans clustering into five clusters (**Fig. 4a**) [37]. This procedure captures the dominant structural variations of each peptide over the simulation period.

**Figure 4.**
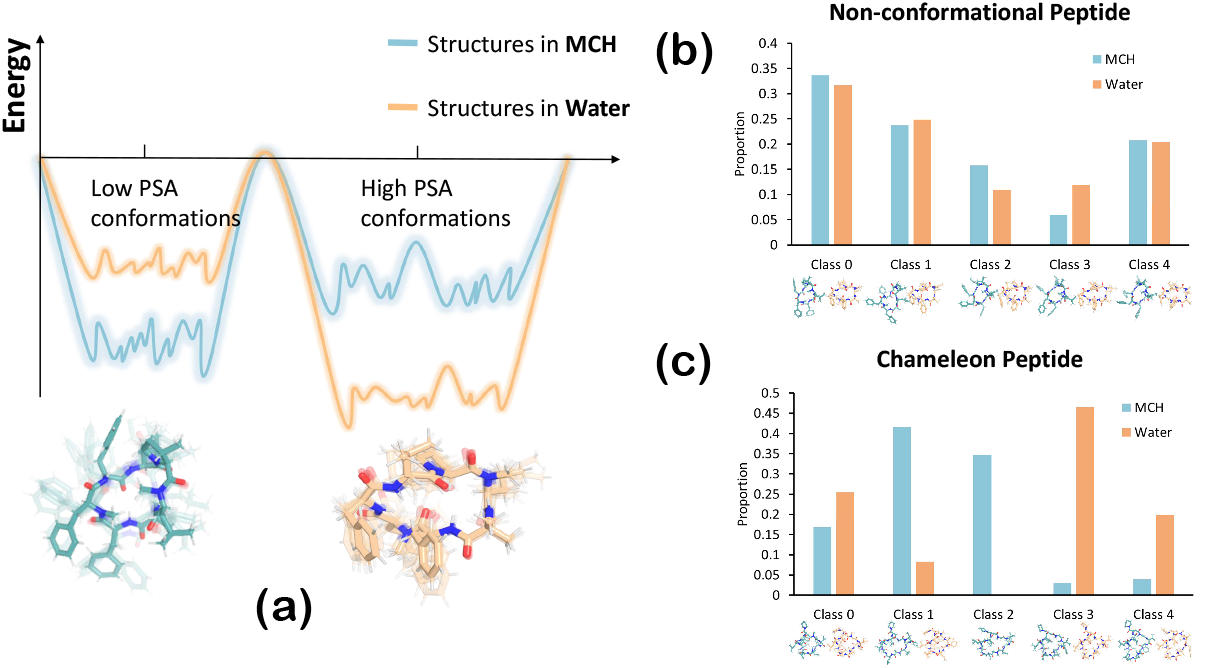
**(a)** Conceptual illustration showing that cyclic peptides can adopt two distinct conformations (high-PSA and low-PSA), whose distributions differ between aqueous and chloroform solvents. For each peptide, 1 *µ*s of MD trajectories (500 ns in water and 500 ns in chloroform at 310 K) were combined and clustered based on backbone atom RMSD, yielding five representative conformational clusters. **(b)** Distribution of clusters for small cyclic peptides with fewer than nine residues, which lack conformational switching and thus exhibit solvent-dependent conformations. **(c)** Distribution of clusters for chameleon cyclic peptides with nine or more residues, which display more similar conformational ensembles across solvents, resulting in smaller ΔPSA values that reflect their adaptive flexibility.

Macrocyclic peptides predominantly adopt two conformational states: a high-PSA ratio state, which exposes hydrogen bond donors and acceptors to the solvent, and a low-PSA ratio state, which favors intramolecular hydrogen bonding. Environmental polarity modulates the population of these states: the high-PSA ratio conformation is favored in aqueous solvent, whereas the low-PSA ratio conformation dominates in chloroform (**Fig. 4a**). Clustering results demonstrate distinct behaviors between small and macrocyclic peptides. Small cyclic peptides (7 residues) exhibit all five clusters with similar proportions in both solvents, confirming the absence of significant conformational switching (**Fig. 4b**). In contrast, macrocyclic peptides (≥9 residues) show substantial solvent-dependent differences in cluster distributions (**Fig. 4c**). For example, Class 2 represents 35% of the population in chloroform but is almost absent in aqueous solvent, whereas Classes 3 and 4 account for 67% of the population in water but are minor species in chloroform. We assign Class 2 to non-polar conformations and Class 3 and 4 to polar conformations. This switching between polar and non-polar conformations underlies the chameleon effect and explains why chameleon peptides display larger Δ*PSA ratio* values.

Together, the analyses in **Fig. 3** and **Fig. 4** demonstrate that ΔPSA serves as a quantitative indicator of conformational adaptability: larger peptides with sufficient flexibility (≥9 residues) can more readily switch between polar and non-polar states across solvent environments, facilitating membrane permeability, whereas smaller peptides remain conformationally restricted. These results provide a rationale for selecting monomer length ≥ 9 as a cutoff for studying chameleon cyclic peptides.

In combination with the analyses presented in **Fig. 3** and **Fig. 4**, macrocyclic peptides are sufficiently long to exhibit chameleon-like conformational adaptability, allowing meaningful evaluation of Δ*PSA ratio* as a metric for conformational switching and membrane permeability.

### 3.2 Benchmark

A set of macrocyclic peptides (monomer length ≥ 9 residues) from the CycPeptMPDB database was subjected to molecular dynamics simulations following a standard protocol (see **Fig. 2a**). The resulting training dataset comprised 2,300 macrocyclic peptides, of which 1,593 (≈ 69.3%) are classified as permeable (permeability *>* –6), and the remaining 707 (≈ 30.7%) as non-permeable (permeability ≤ –6).

For each peptide, 500 ns MD simulations were conducted under four conditions: 310 K in aqueous solvent, 310 K in chloroform solvent, 490 K in aqueous solvent, and 490 K in chloroform solvent, yielding a total simulation time of 2 *µ*s per molecule. All experiments were conducted on a workstation equipped with a 9-core CPU and a high-performance GPU. 500 ns MD simulation for one system with 6.5 k atoms averaged 6 h. Times were recorded via Gromacs. From the trajectories, we computed dynamic and structural features including: SASA, radius of gyration, RMSD, RMSF, number and lifetime of hydrogen bonds, and key bond angles of backbone and side chains. Statistical summaries (mean, median, maximum, minimum, standard deviation, and time-dependent trends) were also extracted, alongside basic molecular descriptors such as total atom count and number of polar/non-polar atoms, resulting in 2,561 features per peptide. These features resulted in the most extensive database to date of macrocyclic peptides containing multiple conformations.

Several machine learning models were trained on our curated dataset for the regression task of predicting cyclic peptide permeability. Among them, the Random Forest model emerged as the top performer (**Table 2**), with an MSE of 0.220, MAE of 0.344, and RMSE of 0.469 [4, 5]. This performance exceeds that of existing models like Multi_CycGT [6] and CycPeptMP [18], demonstrating the superior foundation provided by our dataset for permeability prediction. Based on Random Forest feature importance analysis across five-fold cross-validation (**Table S1**), 35 features were identified as optimal for model stability.

**Table 2.** Performance of Models for Permeability Regression on the Macrocyclic Peptides Dataset with 5-Fold Cross-Validation.

| Model | MAE ↓ | MSE ↓ | RMSE ↓ | R<sup>2</sup> ↑ |
| --- | --- | --- | --- | --- |
| XGBoost | 0.357 ± 0.017 | 0.241 ± 0.023 | 0.490 ± 0.023 | 0.665 ± 0.044 |
| Gradient Boosting | 0.373 ± 0.009 | 0.246 ± 0.012 | 0.496 ± 0.012 | 0.659 ± 0.023 |
| MLP | 0.376 ± 0.014 | 0.252 ± 0.022 | 0.502 ± 0.022 | 0.649 ± 0.041 |
| SVM | 0.361 ± 0.018 | 0.256 ± 0.023 | 0.506 ± 0.023 | 0.642 ± 0.044 |
| CycPeptMP<sup>†</sup> | 0.355 ± 0.007 | 0.253 ± 0.013 | – | **0.772\*** ± 0.011 |
| Multi_CycGT<sup>†</sup> | 0.394 ± 0.020 | 0.269 ± 0.035 | 0.518 ± 0.034 | 0.338 ± 0.090 |
| **Random Forest** | **0.344\*** ± 0.011 | **0.220\*** ± 0.018 | **0.469\*** ± 0.019 | 0.693 ± 0.033 |

↑ and ↓ indicate that higher or lower values are better, respectively. The best value of each metric is indicated in bold, and \* denotes the best metric value statistically superior to the second-best ( $p < 0.05$ )
<sup>†</sup> Metric values were taken from original reports; these datasets differ but are all subsets of CycPeptMPDB

We benchmarked various machine learning models for classifying macrocyclic peptide membrane permeability using our dataset. For this task, the Random Forest classifier achieved the highest F1-score of 0.8248, with multilayer perceptron (MLP) also showing comparable discriminative power (F1-score: 0.8223; see **Table S2**) [8]. To maintain model interpretability and facilitate mechanistic insights, we prioritized models with clear decision boundaries over highly complex deep learning architectures.

Our integrated analysis demonstrates that the Random Forest algorithm is a robust and interpretable model for predicting macrocyclic peptide permeability. It achieved superior performance in both regression (with optimal feature set of 35) and classification tasks, surpassing existing benchmarks and validating the quality of our curated dataset for training effective predictive models.

### 3.3 Explanation and Ablation

#### Sidechain Polarity and Rigidity Govern Membrane Permeability in Macrocyclic Peptides

Using all extracted features, we trained a Random Forest classifier to predict the membrane permeability of macrocyclic peptides [30]. Analysis of feature importance indicated that molecule weight (MW), PSA, radius of gyration (Rg) are the most influential descriptors—consistent with the conclusions established in the previous chapter (**Fig. 5a**). To streamline the model and improve interpretability, we retained only the top five SASA-related features. Here, “water” denotes aqueous trajectories, and “MCH” denotes chloroform trajectories. Notably, the top 5 key descriptors are defined as:

**Figure 5.**
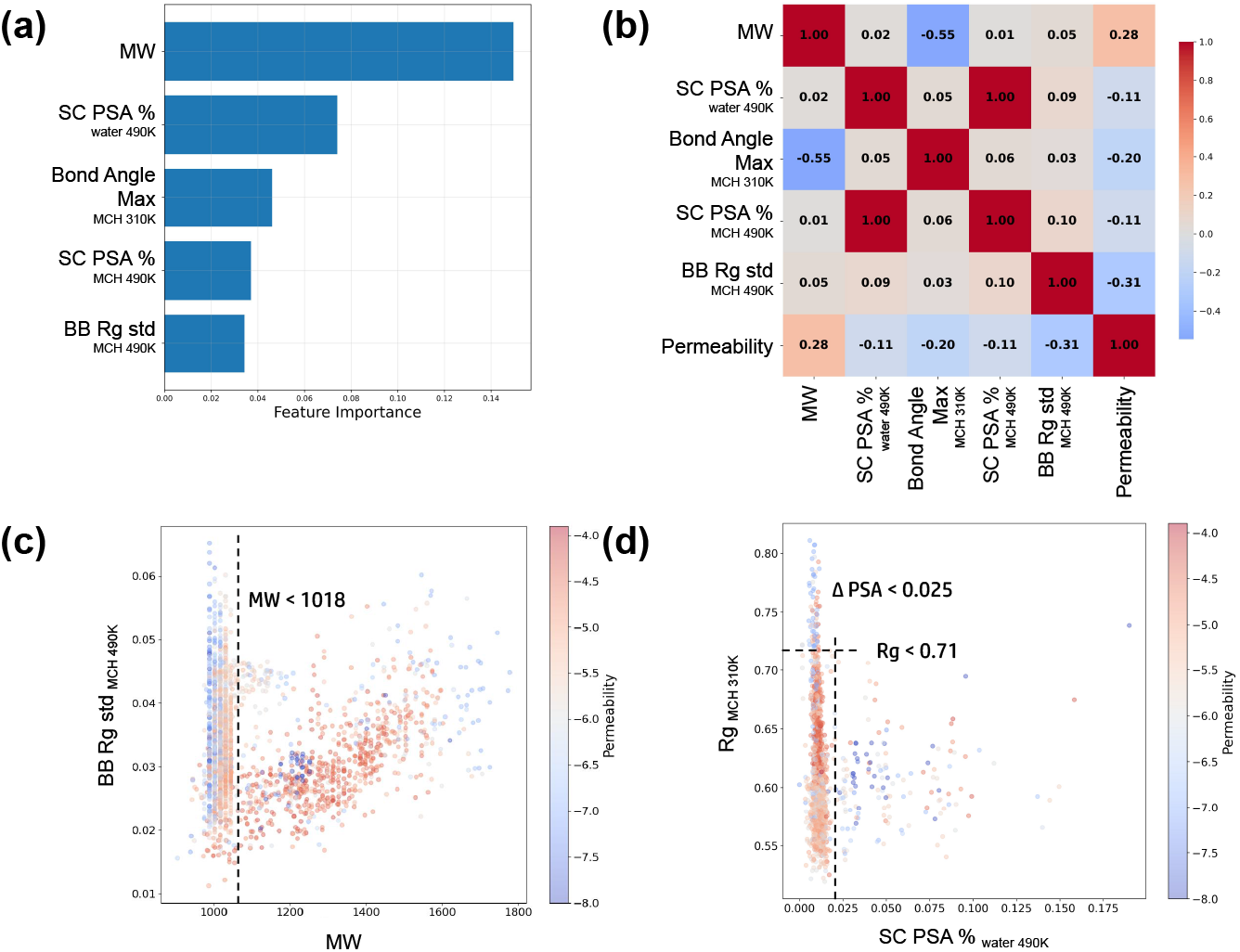
A graphical summary of model interpretability analysis for key predictive features. **(a)** The top five most influential features determined by the Random Forest classifier. **(b)** Correlation relationships between the selected features and the permeability target. **(c)** Data distribution showing how permeability correlates with MW and BB Rg std. **(d)** Data distribution showing how permeability correlates with Rg and SC PSA %.

- *MW* = *Molecule Weight*
- SC PSA 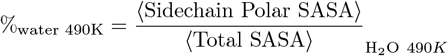
- Bond Angle MaxMCH 310K = max(*Bond Angles*)_CHCl_^3^ 310*K*
- SC PSA 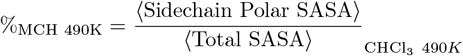
- BB Rg 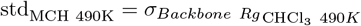 where ⟨·⟩ denotes the average over 500 ns MD simulations at 490 K.

Furthermore, feature correlation analysis revealed that the proportion of sidechain PSA significantly affect the model’s predictions, underscoring the critical role of PSA-related features in determining permeability (**Fig. 5b**) [23]. Based on the analysis of the five key features, an inverse correlation was observed between molecular weight (MW) and membrane permeability, which appears to contradict the conventional wisdom of medicinal chemistry rules such as Lipinski’s Rule of Five. This discrepancy may be attributed to the densely clustered distribution of MW values below 1018 in our dataset, which could introduce anomalous noise and bias into the model (**Fig. 5c**). In the chloroform environment, the standard deviation of backbone radius of gyration (Rg) showed a negative correlation with permeability, suggesting that membrane-permeable macrocyclic peptides exhibit smaller fluctuations in Rg and thus greater conformational stability. This enhanced structural rigidity may indicate stronger intramolecular interactions, as permeable peptides are more likely to form internal hydrogen bonds under chloroform-simulated conditions. Furthermore, membrane-permeable macrocyclic peptides generally exhibit a sidechain polar surface area (SC PSA) proportion below 0.025 in aqueous environments (**Fig. 5d**). This trend implies that higher lipophilicity and more closed conformations of the sidechains contribute to better permeability. However, this distribution may also reflect experimental biases, such as the common practice of incorporating hydrophobic groups (e.g., methylation) to enhance membrane permeability.

**Fig. 5** suggests that these two SASA features alone can effectively capture the conformational dynamics essential for membrane permeability. Highly permeable macrocyclic peptides are characterized by chameleon-like behavior—the ability to reversibly shift between polar and non-polar states. This is evidenced by the retention of discernible non-polar conformations in aqueous environments and a substantial proportion of polar conformations in chloroform. This dual conformational accessibility indicates that the peptides are not kinetically trapped in a single state but can adapt their structure according to environmental polarity—a hallmark of the chameleon effect. Such adaptability allows efficient navigation through the polarity gradient encountered during membrane translocation: polar conformations interact with the aqueous milieu outside the membrane, non-polar conformations traverse the hydrophobic lipid core, and polar conformation

In summary, **Fig. 3** and **Fig. 4** illustrate that macrocyclic peptides (≥ 9 residues) exhibit solvent-dependent conformational changes, reflected in ΔPSA, whereas small cyclic peptides (*<* 9 residues) remain rigid. **Fig. 5** further refines this insight by showing that features capture the key conformational adaptability underlying chameleon-like behavior. Macrocyclic peptides capable of dynamically accessing both polar and non-polar states achieve higher membrane permeability, and these dynamic features provide a concise, quantitative representation of this mechanism.

#### Impact of Simulation Parameters on Molecular Properties

To identify the minimum simulation requirements for robust feature extraction, we ablated our dataset by training Random Forest models on distinct simulation subsets: MCH, water, 490 K, and 310 K. Results in **Table 3** demonstrate that features from the 490 K simulation alone yielded a competitive MSE of 0.2239, matching the performance achieved with the full 500 ns trajectory. Conversely, models relying exclusively on 310 K simulations exhibited markedly reduced predictive efficacy. These findings collectively demonstrate that the 490 K condition alone is sufficient, as it accelerates the system’s convergence to equilibrium, thus efficiently encapsulating the conformational ensemble essential for permeability prediction.

**Table 3.** Model Performance Across Simulation Conditions.

|  | MCH | water | 490K | 310K |
| --- | --- | --- | --- | --- |
| $R^2 \uparrow$ | $0.6863 \pm 0.0277$ | $0.6918 \pm 0.0390$ | $0.6880 \pm 0.0364$ | $0.6053 \pm 0.0390$ |
| MSE ↓ | $0.2254 \pm 0.0137$ | $0.2211 \pm 0.0200$ | $0.2239 \pm 0.0182$ | $0.2838 \pm 0.0232$ |
| RMSE ↓ | $0.4745 \pm 0.0144$ | $0.4697 \pm 0.0212$ | $0.4728 \pm 0.0192$ | $0.5322 \pm 0.0222$ |
| MAE ↓ | $0.3501 \pm 0.0150$ | $0.3467 \pm 0.0169$ | $0.3447 \pm 0.0150$ | $0.3977 \pm 0.0220$ |

Integrating these findings with our earlier analyses (**Fig. 5**), we conclude that the most critical features for predicting macrocyclic peptide membrane permeability are SASA and Rg features at 490 K. According to the convergence analysis, performing two independent 300-ns MD simulations—one in aqueous solvent at 490 K and one in chloroform—can effectively replace longer simulations, substantially reducing computational cost and enabling high-throughput screening.

## 4. Conclusion

In this work, we systematically investigated the relationship between cyclic peptide size, conformational adaptability, and membrane permeability using an integrated approach combining molecular dynamics (MD) simulations and machine learning. Analysis of the CycPeptMPDB dataset revealed that macrocyclic peptides with≥ 9 residues exhibit substantial conformational flexibility, enabling solvent-dependent switching between polar and non-polar states—the so-called ‘chameleon effect’. This behavior is reflected in smaller Δ*PSA* values but larger Δ*PSA ratio*, which serve as a reliable indicator of conformational adaptability and, consequently, membrane permeability.

Through feature extraction based on machine learning, we demonstrated that two features—Rg in MCH and sidechain PSA ratio —serve as robust predictors of macrocyclic peptide permeability. Rg in MCH reflects the peptide’s ability to maintain polar intramolecule interactions in a non-polar, membrane-mimicking environment, while sidechain PSA ratio in water captures the accessibility of hydrophilic surfaces in a polar environment. Together, these two features quantify the peptide’s conformational balance and dynamic adaptability across environments, directly linking structural flexibility to membrane-crossing capability.

Additionally, our study established that two MD simulations under 490K (one in aqueous solvent, one in chloroform) are sufficient for robust feature extraction, drastically reducing computational cost while maintaining predictive performance. This framework provides a high-throughput approach for evaluating cyclic peptide permeability and offers a mechanistic basis for designing chameleon macrocycles with tunable membrane-penetrating properties. While the current study focuses on passive transport driven by conformational changes, it establishes a foundational methodology that can be extended to explore other permeability pathways, such as active transport or the permeation of rigid macrocycles, in future work.

We provide an open-source dataset containing 500-ns MD trajectories conducted in both aqueous and chloroform solvents at both 310K and 490 K. This dataset offers a rich resource for exploring the conformational landscapes of macrocyclic peptides under different solvent conditions, enabling researchers to investigate the relationship between structural dynamics and functional properties such as membrane permeability. By making these data publicly available, we aim to support the broader community in mechanistic studies, model development, and the rational design of chameleon peptides with tailored permeability profiles.

## Supporting Information

The Supporting Information is available free of charge at \href{url}{description}.

**Figure S1**

Distribution Analysis of CycPeptMPDB

**Figure S2**

Different Δ*SASA*^*∗*^ distribution with permeability

**Table S1**

Five-Fold Cross-Validation Results for Permeability Regression Using Machine Learning Models

**Table S2**

Five-Fold Cross-Validation Results for Permeability Classification Using Machine Learning Models

**Table S3**

Model Performance Across Different Simulation Conditions

## Acknowledgments

We thank just about everybody.

