## Supporting Information for "Delta PSA: A New Metric for Conformational Dynamics Underlying Macrocyclic Peptide Permeability"

#### Figures

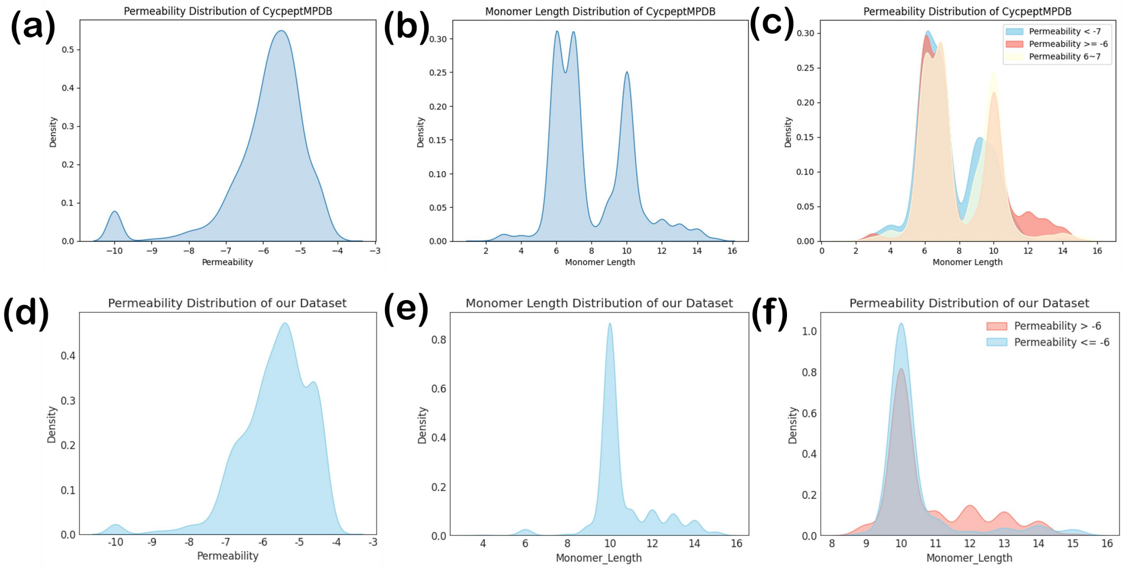


**Figure S1**

(a)-(c) Distributions of Permeability and Monomer Length for all samples in the CycPeptMPDB database.

(d)-(f) Distributions of Permeability and Monomer Length for samples in our selected dataset.

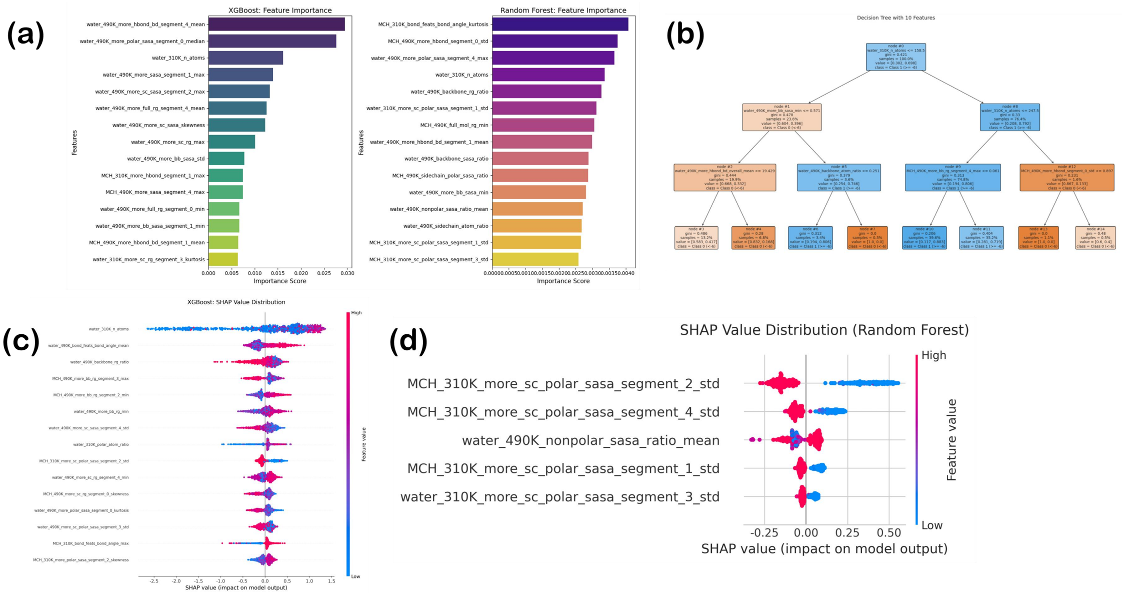


**Figure S2**

(a) Feature importance plots for Random Forest and XGBoost models in the regression task, showing the top 15 features selected based on the feature importance scores provided by each model.

(b) A simplified decision tree model derived from the Random Forest regression model. This tree visually illustrates the key decision rules and feature splits used by the Random Forest to make predictions.

(c)-(d) SHAP value plots for the Random Forest and XGBoost regression models, respectively. These plots demonstrate the influence of each feature on individual predictions by showing the SHAP values, which quantify the contribution of each feature to pushing the model output away from the base value. Positive SHAP values indicate a feature increases the predicted value, while negative values indicate a decrease, providing insights into how specific features affect the regression results.

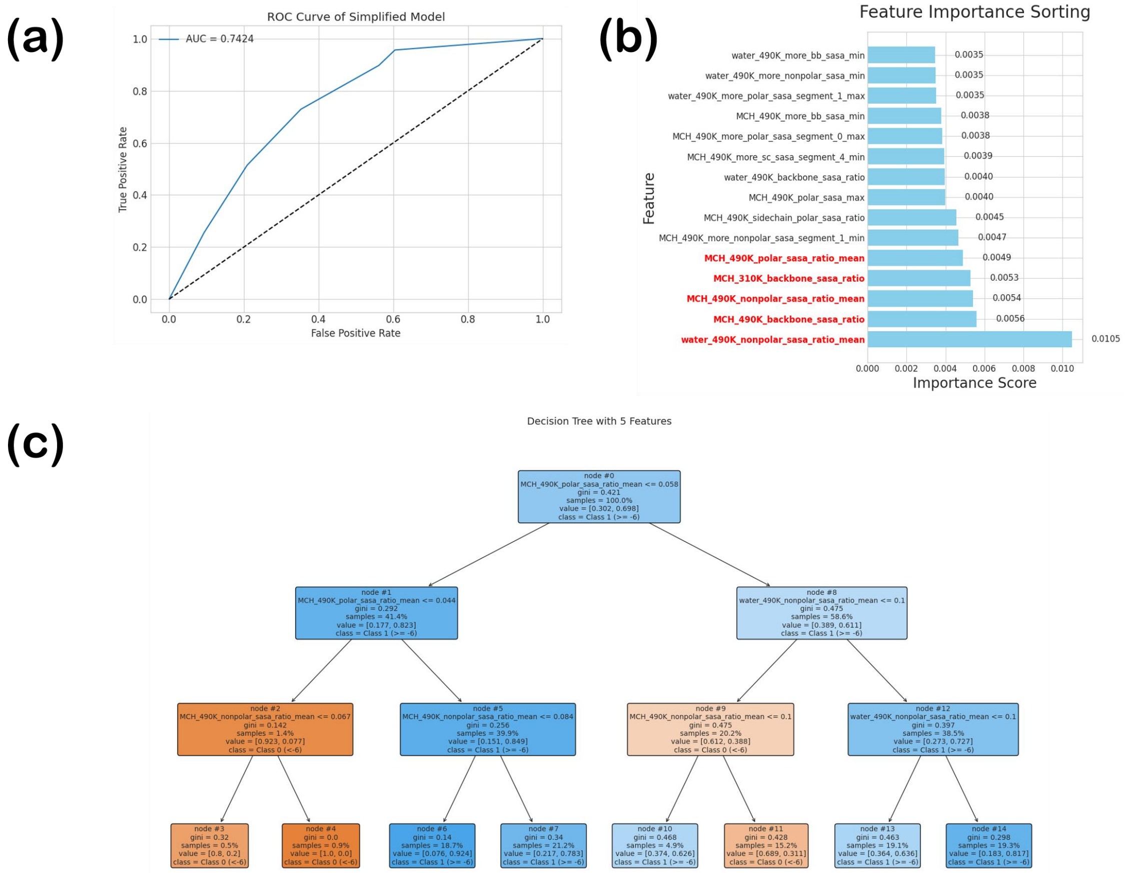


**Figure S3**

(a) AUC-ROC curve of the Random Forest discriminant model trained on the top 5 SASA-related features.

(b) Feature ranking based on the internal feature importance scores of the model. This ranking reflects the relative contribution of each of the top 5 SASA-related features to the model's decision-making process.

(c) Simplified decision tree derived from the Random Forest discriminant model.

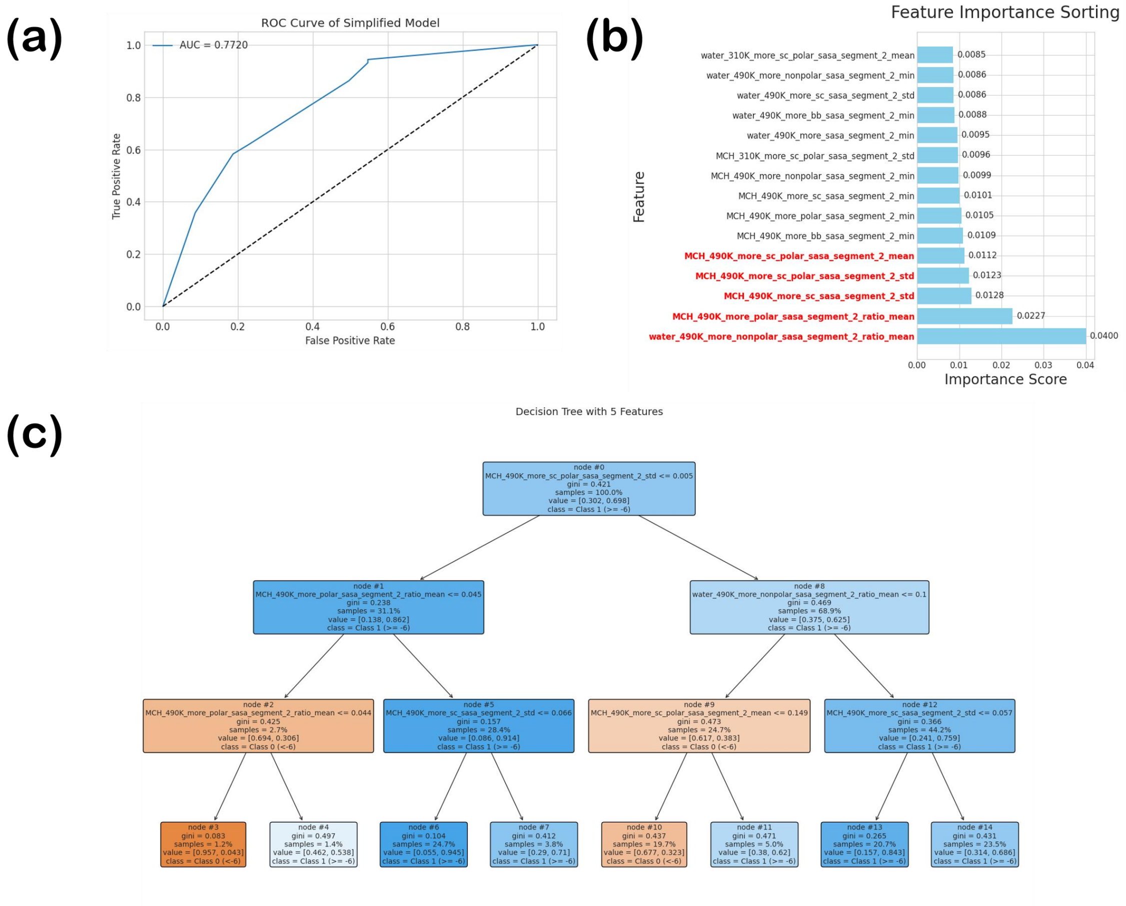


**Figure S4**

(a) AUC-ROC curve of the Random Forest discriminant model trained on the top 5 SASA-related features extracted from 0–300 ns trajectories.

(b) Feature ranking based on the internal feature importance scores of the model. This ranking reflects the relative contribution of each of the top 5 SASA-related features ( extracted from 0–300 ns trajectories ) to the model's decision-making process.

(c) Simplified decision tree derived from the Random Forest discriminant model.

### Tables

**Table S1.** Performance of machine learning models on the permeability regression task across each fold in five-fold cross-validation, including Mean Absolute Error (MAE), Mean Squared Error (MSE), Root Mean Squared Error (RMSE), and R-squared ($R^2$) for all folds.

| Model | fold_num | MAE | MSE | RMSE | $R^2$ |
|---|---|---|---|---|---|
| SVM | 1 | 0.4316 | 0.3373 | 0.5808 | 0.5375 |
| | 2 | 0.4267 | 0.3136 | 0.5600 | 0.5745 |

| | | | | | |
|---|---|---|---|---|---|
| | 3 | 0.4152 | 0.3338 | 0.5778 | 0.5241 |
| | 4 | 0.4312 | 0.3268 | 0.5717 | 0.5397 |
| | 5 | 0.4050 | 0.3253 | 0.5704 | 0.5531 |
| Random Forest | 1 | 0.3910 | 0.2701 | 0.5198 | 0.6296 |
| | 2 | 0.3968 | 0.2718 | 0.5214 | 0.6312 |
| | 3 | 0.3901 | 0.3003 | 0.5480 | 0.5719 |
| | 4 | 0.3923 | 0.2649 | 0.5147 | 0.6269 |
| | 5 | 0.3892 | 0.2838 | 0.5328 | 0.6101 |
| XGBoost | 1 | 0.4173 | 0.3034 | 0.5508 | 0.5840 |
| | 2 | 0.3907 | 0.2786 | 0.5278 | 0.6220 |
| | 3 | 0.4090 | 0.3119 | 0.5585 | 0.5554 |
| | 4 | 0.4056 | 0.2965 | 0.5446 | 0.5824 |
| | 5 | 0.3929 | 0.2919 | 0.5403 | 0.5990 |
| KNN | 1 | 0.4414 | 0.3516 | 0.5929 | 0.5179 |
| | 2 | 0.4261 | 0.3425 | 0.5852 | 0.5353 |
| | 3 | 0.4314 | 0.3671 | 0.6059 | 0.4766 |
| | 4 | 0.4439 | 0.3479 | 0.5898 | 0.5101 |
| | 5 | 0.4318 | 0.3809 | 0.6172 | 0.4767 |

**Table S2**. Performance of machine learning models on the permeability classification task across each fold in five-fold cross-validation, including Mean Absolute Error (MAE), Mean Squared Error (MSE), Root Mean Squared Error (RMSE), and R-squared ($R^2$) for all folds.

| Model | fold_num | accuracy | precision | f1_score | recall | auc | mcc |
|---|---|---|---|---|---|---|---|
| SVM | 1 | 0.789130 | 0.814607 | 0.856721 | 0.903427 | 0.829815 | 0.470519 |
| | 2 | 0.806522 | 0.806878 | 0.872675 | 0.950156 | 0.858054 | 0.509884 |
| | 3 | 0.795652 | 0.797900 | 0.866097 | 0.947040 | 0.848204 | 0.478596 |
| | 4 | 0.821739 | 0.823848 | 0.881159 | 0.947040 | 0.867366 | 0.552634 |
| | 5 | 0.795652 | 0.811475 | 0.863372 | 0.922360 | 0.824725 | 0.480005 |
| XGBoost | 1 | 0.819565 | 0.836158 | 0.877037 | 0.922118 | 0.856586 | 0.550516 |
| | 2 | 0.802174 | 0.830460 | 0.863976 | 0.900312 | 0.869898 | 0.509134 |
| | 3 | 0.836957 | 0.839779 | 0.890190 | 0.947040 | 0.875614 | 0.594133 |
| | 4 | 0.815217 | 0.837143 | 0.873323 | 0.912773 | 0.879827 | 0.541177 |
| | 5 | 0.789130 | 0.816901 | 0.856721 | 0.900621 | 0.822216 | 0.469061 |
| RandomForest | 1 | 0.797826 | 0.822034 | 0.862222 | 0.906542 | 0.850232 | 0.494306 |
| | 2 | 0.802174 | 0.817680 | 0.866764 | 0.922118 | 0.855095 | 0.501637 |
| | 3 | 0.806522 | 0.815217 | 0.870827 | 0.934579 | 0.863108 | 0.511286 |
| | 4 | 0.830435 | 0.840336 | 0.884956 | 0.934579 | 0.876499 | 0.577775 |
| | 5 | 0.795652 | 0.822034 | 0.860947 | 0.903727 | 0.863950 | 0.486653 |
| KNN | 1 | 0.786957 | 0.810585 | 0.855882 | 0.906542 | 0.821533 | 0.462951 |

| | | | | | | | |
|---|---|---|---|---|---|---|---|
| | 2 | 0.752174 | 0.777480 | 0.835735 | 0.903427 | 0.794460 | 0.359169 |
| | 3 | 0.776087 | 0.794595 | 0.850941 | 0.915888 | 0.801912 | 0.427279 |
| | 4 | 0.797826 | 0.831395 | 0.860150 | 0.890966 | 0.821847 | 0.500904 |
| | 5 | 0.743478 | 0.796512 | 0.822823 | 0.850932 | 0.754973 | 0.362677 |

**Table S3. Performance of different features on the permeability classification task.** To evaluate the contribution of selected features, the performance of the Random Forest model with simplified feature sets is reported.

| Model with Simplified Features | accuracy | precision | f1_score | Recall | Auc | Mcc |
|---|---|---|---|---|---|---|
| Simplified_with_5_features | 0.7500 | 0.8160 | 0.8223 | 0.8287 | 0.7563 | 0.4012 |
| Simplified_with_10_features | 0.7935 | 0.8139 | 0.8605 | 0.9128 | 0.7614 | 0.4796 |
| Simplified_with_5_sasa | 0.7870 | 0.7852 | **0.8624** | 0.9564 | 0.7424 | **0.4528** |
| Simplified_with_10_sasa | 0.7674 | 0.7675 | 0.8516 | 0.9564 | 0.7121 | 0.3918 |
